# Suppression of p16 alleviates the senescence-associated secretory phenotype

**DOI:** 10.1101/2020.08.19.257717

**Authors:** Raquel Buj, Kelly E. Leon, Katherine M. Aird

## Abstract

Oncogene induced senescence (OIS) is characterized by increased expression of the cell cycle inhibitor p16, leading to a hallmark cell cycle arrest. Suppression of p16 in this context drives proliferation, senescence bypass, and contributes to tumorigenesis. OIS cells are also characterized by the expression and secretion of a widely variable group of factors collectively termed the senescence-associated secretory phenotype (SASP). The SASP can be both beneficial and detrimental and affects the microenvironment in a highly context-dependent manner. The relationship between p16 suppression and the SASP remains unclear. Here, we show that knockdown of p16 decreases expression of the SASP factors and pro-inflammatory cytokines *IL6* and *CXCL8* in both RAS^G12V^-and BRAF^V600E^-induced senescence. Notably, this is likely not due to suppression of senescence as *LMNB1* and cyclin A expression remain low and p21 remains high. Moreover, low p16 expression in both cancer cell lines and patient samples correspond to decreased SASP gene expression, suggesting this is a universal effect of loss of p16 expression. Together, our data suggest that p16 transcriptionally regulates SASP gene expression, which has implications for understanding how p16 modulates both the senescent and tumor microenvironment.

## Introduction

Senescence is considered a state of stable cell cycle arrest that can occur due to a variety of stimuli [1]. Oncogene-induced senescence (OIS) occurs upon activation of an oncogene such as HRAS or BRAF in normal cells [2, 3]. One of the hallmarks of senescent cells is upregulation of the cell cycle inhibitor *CDKN2A* (encoding for p16), which restrains cell cycle progression and cellular proliferation [4–6]. Canonically, elevated p16 represses hyperphosphorylation of the retinoblastoma protein (RB), which inhibits E2F transcription factor-mediated expression of proliferative genes [7]. Loss of p16 is a common event in human cancer that has been linked to senescence bypass, increased proliferation, and malignant transformation though both canonical and non-canonical (RB-independent) pathways [8–12].

The acquisition of a senescence-associated secretory phenotype (SASP) is also characteristic of OIS cells [13]. The SASP is composed of a variety of soluble signaling factors including pro-inflammatory cytokines, chemokines, and growth factors, as well as proteases, insoluble extracellular matrix proteins and non-protein components that are transcriptionally and translationally upregulated and secreted into the surrounding microenvironment by senescent cells [14–18]. Due to the impact that SASP can exert on cellular physiology, this program is tightly regulated at multiple levels. At the transcriptional level, several transcription factors (NF-κB, C/EBP-β) and upstream regulators (p38 MAPK, GATA4, p53, and ATM) have been described to either positively or negatively regulate SASP gene expression [16, 19–25]. The SASP is also regulated at both the epigenetic [26–31] and translational level [17, 32]. Recent publications suggest that the initiation of SASP gene transcription during OIS is likely due to loss of lamin B1 (*LMNB1*) and nuclear integrity [33, 34], leading to the accumulation of cytoplasmic chromatin fragments (CCFs) [35, 36]. CCFs activate the cytosolic DNA sensor cyclic guanosine monophosphate (GMP)-adenosine monophosphate (AMP) synthase (cGAS) that catalyzes the synthesis of the second messenger cyclic GMP-AMP (cGAMP) to bind and activate stimulator of interferon genes (STING), leading to NF-κB activation and cytokine transcription [35, 37, 38]. Therefore, changes in LMNB1 expression are tightly linked to SASP gene transcription.

It is well documented that the SASP modifies the cellular microenvironment and alters neighboring cells, exerting a pleiotropic effect that is not fully understood [39]. On one hand, SASP factors contribute to wound-healing [40–42], normal development [43, 44], and have tumor suppressive effects through the recruitment of different immune cells to clear premalignant cells, a process termed senescence surveillance [45–47]. On the other hand, SASP factors can be pro-tumorigenic by sustaining proliferation, invasion, metastasis, and chemoresistance [48–52]. This paradoxical double role of SASP factors is highly dependent on both genetic background and SASP composition, which is known to be both variable and dynamic [53]. Different genetic backgrounds, cellular contexts, and/or senescence inducers allow for different SASP programs that can promote or inhibit tumorigenesis [54–56]. Interestingly, different SASP programs can also induce senescence in neighboring cells in a paracrine manner that in turn express a particular SASP program [54]. Thus, the final beneficial or detrimental outcome net effect of the SASP is governed by multiple mechanisms that are not yet fully understood [57]. Characterizing whether different genetic backgrounds lead to different SASP programs may be critical to develop efficient and personalized regimens for cancer patients. As ~50% of all human tumors have low p16 expression [58], understanding its role in regulating the SASP has implications for a large subset of patients.

Here, we investigated the effect of p16 suppression on SASP gene expression. We found that knockdown of p16 leads to decreased *IL6* and *CXCL8* (encoding IL8) SASP gene expression in both HRAS^G12V^ and BRAF^V600E^ models of OIS. This was not due to increased *LMNB1* expression, indicating that these changes were not simply an artifact of reduced senescence or inhibited upstream signaling. We confirmed these results in p16-wildtype melanoma cells upon knockdown of p16. Moreover, using publicly-available data, we found that low *CDKN2A* expression in patient tumors is associated with a tumor-characteristic decrease in specific SASP programs. Together, our results suggest that p16 may have a role in transcriptionally regulating SASP factors, which has implications for understanding how loss of p16 affects the senescent and tumor microenvironment.

## Results

### Knockdown of p16 abrogates oncogene-induced *IL6* and *CXCL8* expression

Upregulation of both p16 and SASP factors are characteristic of OIS cells [13]. A previous study found that overexpression of p16 induces senescence without upregulation of the SASP [59]. However, it is unknown whether p16 upregulation is necessary for SASP gene expression in the context of OIS. In order to better understand the effects of p16 expression on the SASP, we assessed the expression of the most extensively characterized interleukins upregulated in senescence, IL6 and IL8 [13, 16, 60]. Concomitant knockdown of p16 with BRAF^V600E^ or HRAS^G12V^ overexpression decreased *IL6* and *CXCL8* expression in IMR90 fibroblasts (Fig. 1A-E), a classical model of OIS [5, 61]. The shRNA hairpin specificity targeting p16 was previously validated [8]. Similar results were observed in normal skin fibroblasts Hs 895.Sk (Fig. 1F-G), suggesting this is not a cell line-specific phenomenon. Together, these data demonstrate that knockdown of p16 abrogates *IL6* and *CXCL8* expression due to oncogene activation.

**Figure 1.**
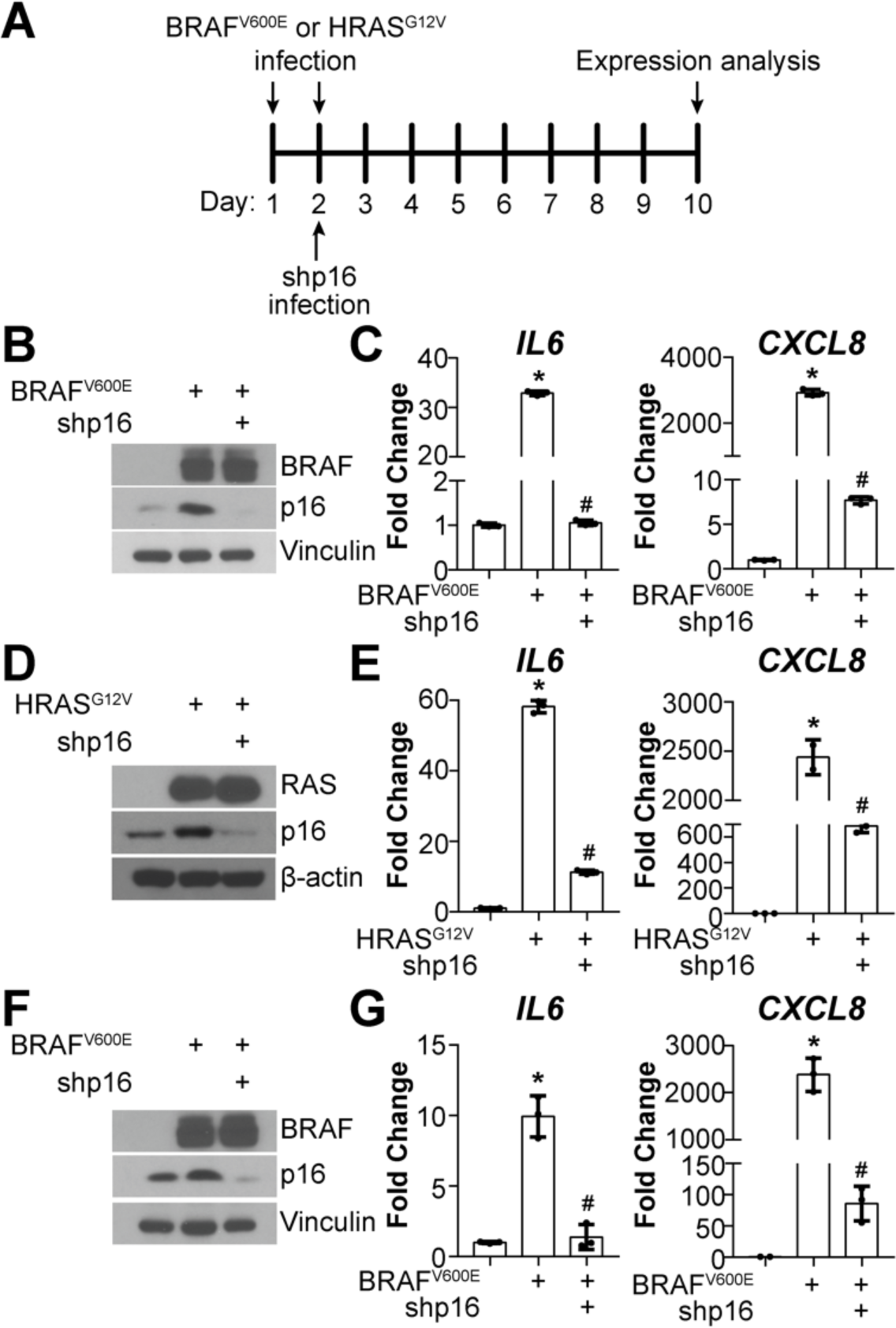
Knockdown of p16 decreases *IL6* and *CXCL8* expression in OIS cells. **(A)** Schematic of infection of cells. **(B-C)** IMR90 cells expressing BRAF^V600E^ alone or in combination with shRNA targeting p16 (shp16). **(B)** Immunoblot of the indicated proteins. **(C)** *IL6* and *CXCL8* mRNA expression. **(D-E)** IMR90 cells expressing HRAS^G12V^ alone or in combination with shp16. **(D)** Immunoblot of the indicated proteins. **(E)** *IL6* and *CXCL8* mRNA expression. **(F-G)** Hs 895.SK cells expressing BRAF^V600E^ alone or in combination with shp16. **(F)** Immunoblot of the indicated proteins. **(G)** *IL6* and *CXCL8* mRNA expression. Expression data show fold change relative to control mean +/-SD. Expression of target genes was normalized against multiple reference genes. Data normalized against *MRPL9* is represented. *p<0.05 vs. control; #p<0.05 vs. BRAF^V600E^ or HRAS^G12V^ alone.

We previously published that knockdown of p16 concomitantly with expression of BRAF^V600E^ bypasses senescence in IMR90 cells [8]. Therefore, it is possible that the SASP expression is low because the cells never undergo OIS. To investigate whether the observed decrease in *IL6* and *CXCL8* is a direct effect of p16 suppression and not simply a consequence of senescence bypass, we knocked down p16 at different time points after oncogene expression (Fig. 2A). Consistent with our previous data, knockdown of p16 at these different timepoints also decreased both *IL6* and *CXCL8* expression (Fig. 2B-D). Notably, this was not due to a decrease in senescence as p21 remained high and cyclin A and *LMNB1* expression remained low (Fig. 2B & 2E). These data suggest that p16 regulates *IL6* and *CXCL8* expression, which is uncoupled from senescence bypass.

**Figure 2.**
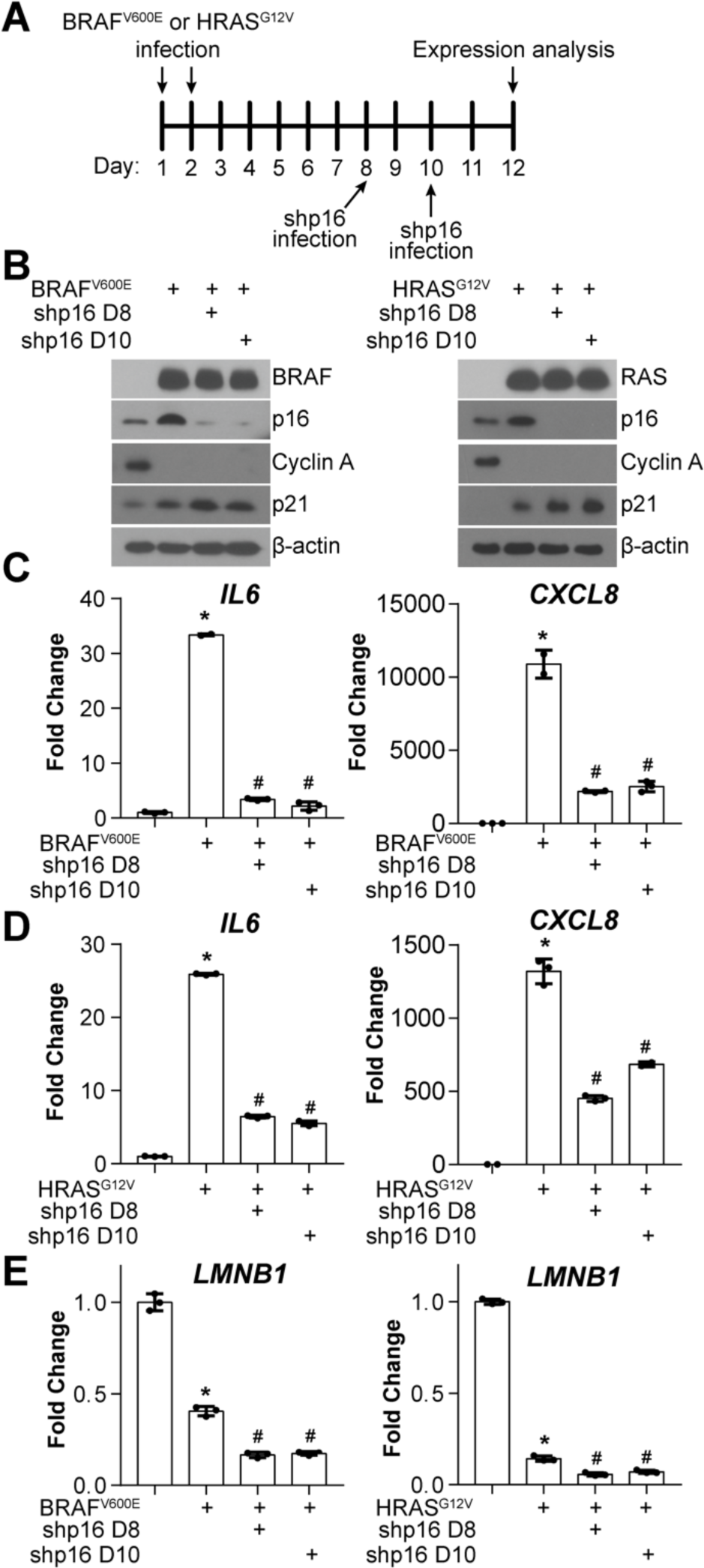
Knockdown of p16 at later timepoints decreases *IL6* and *CXCL8*, while not increasing *LMNB1* expression in OIS cells. **(A)** Schematic of infection of cells.**(B-E)** IMR90 cells expressing BRAF^V600E^ or HRAS^G12V^ alone or in combination with shRNA targeting p16 (shp16) at the indicated time points (D8 = day 8; D10 = day 10). **(B)** Immunoblot of the indicated proteins. **(C)** *IL6* and *CXCL8* mRNA expression in the BRAF^V600E^ model. **(D)** *IL6* and *CXCL8* mRNA expression in the HRAS^G12V^ model. **(E)** *LMNB1* mRNA expression. Expression data show fold change relative to control mean +/-SD. Expression of target genes was normalized against multiple reference genes. Data normalized against *MRPL19* is represented. *p<0.05 vs. control; #p<0.05 vs. BRAF^V600E^ or HRAS^G12V^ alone.

### Knockdown of p16 in tumor cells decreases *IL6* and *CXCL8* expression

Next, we aimed to investigate whether suppression of p16 leads to decreased expression of *IL6* and *CXCL8* in tumor cells. Towards this goal, we knocked down p16 in three melanoma cell lines with wildtype p16 (Fig. 3A). Similar to normal fibroblasts, knockdown of p16 decreased both *IL6* and *CXCL8* in the melanoma cells (Fig. 3B-D). Consistent with our data in fibroblasts (Fig. 2), suppression of p16 in the melanoma cells also decreased *IL6* and *CXCL8* expression. Altogether, these data suggest that p16 transcriptionally regulates both *IL6* and *CXCL8* and support the hypothesis that this is not a consequence of p16 suppression-mediated senescence bypass since suppression of p16 in both OIS and proliferating melanoma cells abrogates *IL6* and *CXCL8*.

**Figure 3.**
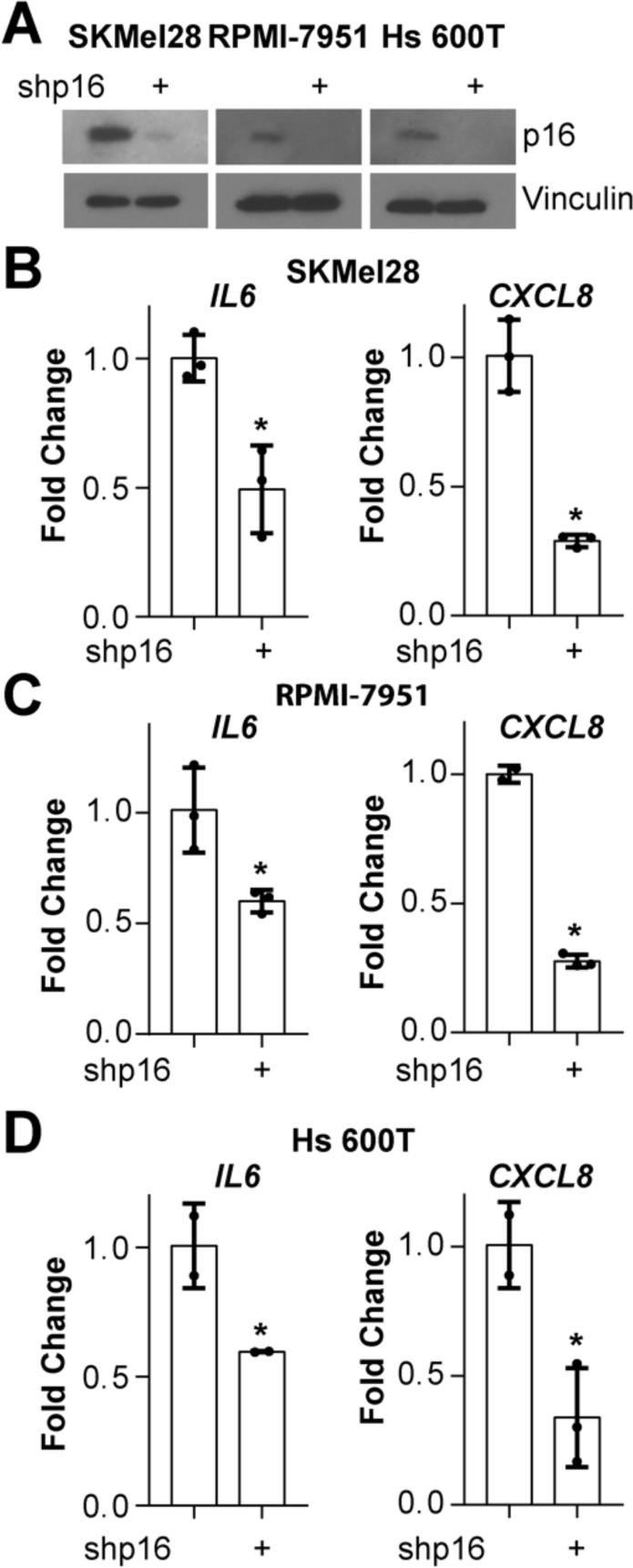
Knockdown of p16 in melanoma cells decreases *IL6* and CXCL8 expression. The melanoma cell lines SKMel28, RPMI-7951, and Hs 600T expressing wildtype p16 were infected with lentivirus expressing shRNA targeting p16 (shp16). **(A)** Immunoblot of the indicated proteins.**(B)** mRNA expression of *IL6* and *CXCL8* in SKMel28 melanoma cells. **(C)** mRNA expression of *IL6* and *CXCL8* in RPMI-7951 melanoma cell lines. **(D)** mRNA expression of *IL6* and *CXCL8* in Hs 600T melanoma cell lines. Expression data show fold change relative to control mean +/-SD. Expression of target genes was normalized against multiple reference genes. Data normalized against *MRPL9* is represented. *p<0.05.

### Low *CDKN2A* in patient tumors correlates with low SASP expression

It has been widely demonstrated that suppression of p16 leads to increased proliferation, tumorigenesis, and metastasis *in vitro* and *in vivo*, and loss of p16 expression is considered a poor prognostic maker [62–65]. To further understand the relationship between loss of p16 and decreased expression of the SASP, we used TCGA data from skin cutaneous melanoma (SKCM, n=473), pancreatic adenocarcinoma (PAAD, n=183), and colorectal adenocarcinoma (COADREAD, n=82), which are all in part characterized by loss of p16 expression [66]. Patients were classified according to their *CDKN2A* status (p16 low or high, see Methods for details) (Table 1), and differential expression analysis was performed independently for each tumor type. Consistently with our previous findings, most of the SASP factors profiled upon RAS-induced senescence (including soluble factors and exosomes, 232 total unique genes) (**Table S1**) [53] were significantly downregulated in p16-low samples in all three tumor types (Fig. 4A and **Table S2**). Interestingly, only 4 SASP factors (*HLA-A*, *TNIP2*, *ACTB*, and *TTYH3*) were significantly downregulated in all 3 tumor types (Fig. 4B and **Table S3**), suggesting that although decreased expression of SASP factors is a characteristic of p16-low tumors, each tumor type displays a distinct profile. Consistent with this observation, Gene Set Enrichment Analysis (GSEA) revealed that pathways related to the SASP, inflammation, and the immune system such as: ‘Senescence Associated Secretory Phenotype’, “Antigen Processing Cross Presentation”, and “Cytosolic DNA Sensing Pathway” where negatively enriched (i.e., negative Normalized Enrichment Score, NES) in p16-low samples in all three tumor types (Fig. 4C and **Table S4**). These data suggest that p16 at least in part transcriptionally regulates SASP factors in human tumor samples. Finally, since we observed downregulation of the “Cytosolic DNA Sensing Pathway” signature, we correlated the expression level of *CDKN2A* and *LMNB1* in the tumors. We did not observe a strong correlation between *CDKN2A* and *LMNB1* expression (Fig. 4D), suggesting that p16 regulation of this pathway and the SASP in tumors is not through modulation of *LMNB1*. All together with our *in vitro* data, these data in human tumor samples demonstrate a universal, positive correlation between p16 expression and SASP gene expression.

**Table 1.** Statistics of TCGA data sets

|  | **SKCM** | **PAAD** | **COADREAD** |
| --- | --- | --- | --- |
| **Total cases** | 473 | 183 | 82 |
| ***CDKN2A* expression (RSEM) mean** | 560.20 | 359.17 | 332.09 |
| ***CDKN2A* expression (RSEM) standard deviation (SD)** | 204.21 | 84.31 | 677.87 |
| ***CDKN2A* expression (RSEM) median** | 1103.44 | 584.37 | 113.29 |
| ***CDKN2A* expression (RSEM) first quartile ($Q_1$)** | 20.21 | 31.92 | 64.39 |
| ***CDKN2A* expression (RSEM) third quartile ($Q_3$)** | 638.01 | 340.72 | 314.50 |
| **No. p16-low cases (*CDKN2A* expression $\leq Q_1$)** | 119 | 46 | 21 |
| **No. p16-high cases (*CDKN2A* expression $\geq Q_3$)** | 119 | 46 | 21 |

**Figure 4.**
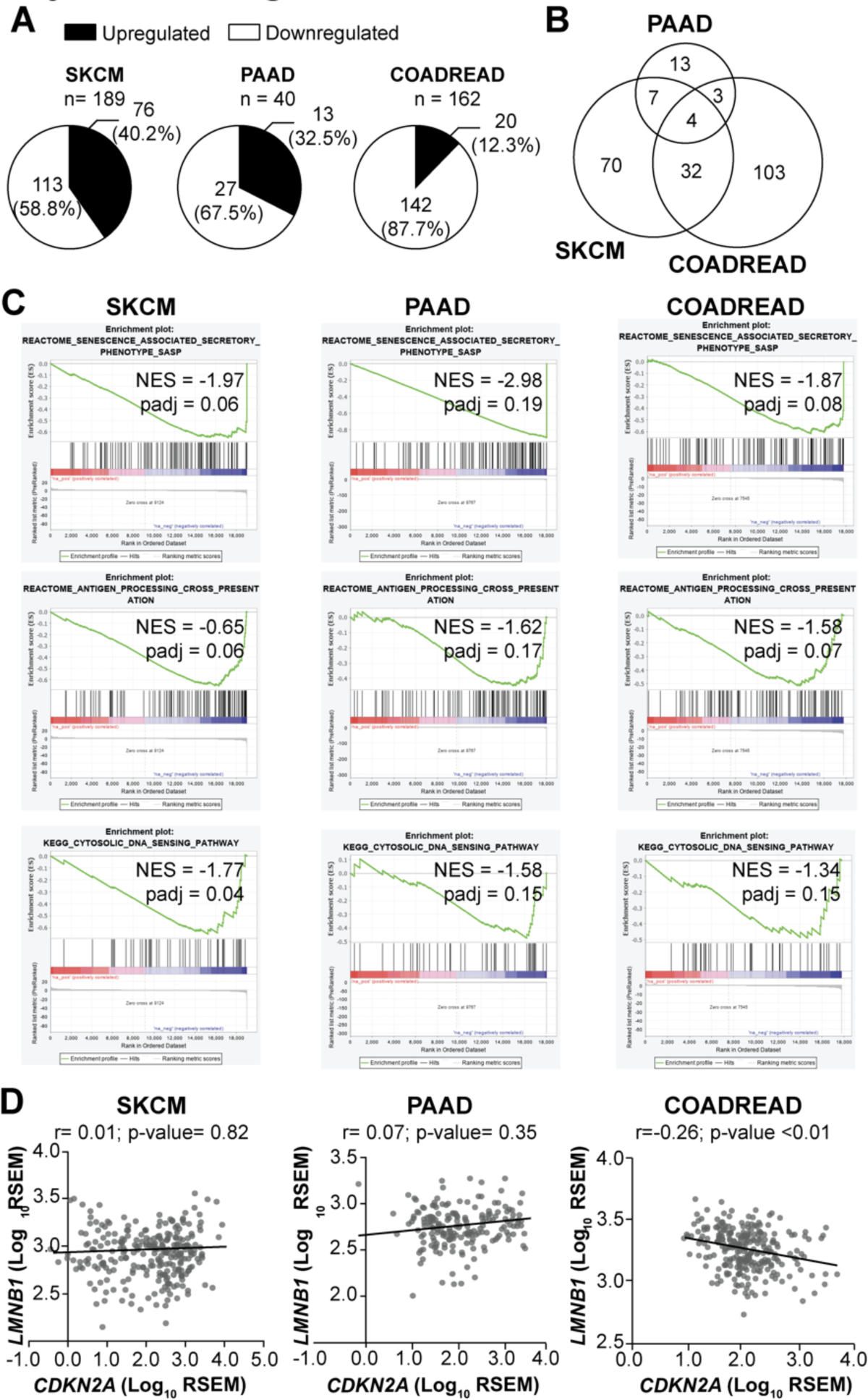
Tumors with low *CDKN2A* expression have decreased expression of SASP and inflammation-related signatures. **(A)** Percentage of SASP genes significantly upregulated and downregulated in *CDKN2A*-low (i.e., p16-low) expressing tumors when compared to *CDKN2A*-high (p16-high) expressing tumors. **(B)** Cross-comparison of significantly downregulated SASP genes in *CDKN2A*-low expressing tumors vs. *CDKN2A*-high expressing tumors in the three tumor types. **(C)** SASP and inflammation-related common negatively enriched terms among the three studied tumor types in Gene Set Enrichment Analysis (GSEA) between *CDKN2A*-low and *CDKN2A*-high expressing tumors. SKCM (skin cutaneous melanoma), PAAD (pancreatic adenocarcinoma), COADREAD (colorectal adenocarcinoma), NES (negative enrichment score). **(D)** Correlation between *CDKN2A* and *LMNB1* expression for each tumor type. Data are shown as Log_10_ of RSEM. Coefficient of correlation (R) and p-value were calculated using Pearson correlation.

## Discussion

Increased expression of p16 and SASP are characteristics of OIS; however, the relationship between them is not well understood. While increased p16 expression has a clear role in sustaining the characteristic cell cycle arrest of OIS cells [8, 67], the SASP appears to be a consequence of the DNA damage response induced by OIS and is not necessary for the proliferative arrest [24, 68, 69]. Indeed, this is evident by the observation that p16-mediated senescence induction, a stimulus that a priori only restrains cell cycle progression, does not induce SASP expression [59]. Here we found that knockdown p16 decreases gene expression of two of the most well characterized SASP factors *IL6* and *CXCL8* [13, 16, 60], while there was no concurrent increase in *LMNB1* and cyclin A nor decreased p21 expression (Fig. 1 & 2). Together, this suggests that p16 expression is not sufficient but is necessary to induce transcription of the SASP.

One important question that remains is how does p16 loss mechanistically affect SASP gene expression? p16 is a critical cell cycle regulator, and its suppression both enhances proliferation and allows for senescence bypass [8–10, 12, 70, 71]. Decreased *LMNB1* expression plays an important role in the establishment of the SASP [72]. Recent publications demonstrate that decreased expression of *LMNB1* and consequential decrease of nuclear integrity leads to the accumulation of CCFs that in turn activate the cGAS-STING signaling pathway to drive the SASP [35–37, 73, 74]. Importantly, we found that knockdown of p16 decreased SASP gene expression, which was not a consequence of increased *LMNB1* expression (Fig. 2). Therefore, this is likely not due to downregulation of cGAS-STING signaling, which is directly affected by LMNB1 and nuclear integrity [35]. Notably, we found that patient tumors with low p16 expression have decreased cytosolic DNA sensing pathway signaling in comparison with p16 high tumors (Fig. 4C); however, no correlation was found between *CDKN2A* and *LMNB1* mRNA expression (Fig. 4D), suggesting that additional mechanisms are at play in tumors with low p16 expression. Loss of *CDKN2A* is often due to deletion or hypermethylation of the locus [66]. Interestingly, previous work has suggested that melanomas with low MTAP have decreased cGAS-STING signaling [75], and *MTAP* is often co-deleted/silenced with *CDKN2A* [76]. Therefore, multiple mechanisms may exist in tumors with loss of this locus to suppress SASP gene expression and/or modulate the tumor microenvironment. Additionally, p16 can negatively regulate TP53 (encoding for p53) at the transcriptional level and also at the protein level by increasing Mdm2-dependent degradation of p53 [77, 78]. As p53 is a negative regulator of the SASP [79], it is possible that the observed decrease in SASP expression upon p16 suppression is due to negative regulation of p53. Future studies are needed to determine the exact mechanism by which p16 suppression decreases SASP gene expression.

We and others have shown that in addition to its canonical role regulating cell cycle progression though the RB pathway, p16 has non-canonical activities that regulate other important aspects of cellular physiology such as nucleotide metabolism, reactive oxygen species, and miRNAs among others [80, 81]. In this regard, some studies have found that pharmacological inhibition of canonical downstream targets of p16, namely CDK4/6, leads to an induction of SASP factors, recruitment of antitumor immune cells, and senescence [82–85], suggesting that p16 regulation of SASP expression and the tumor immune microenvironment may be due to non-canonical (RB-independent) mechanisms. On the contrary, other authors suggest that inhibition of CDK4/6 alone does not induce a SASP and immunologic responses [86]. Additional studies are needed to delineate the exact mechanism whereby suppression of p16 decreases SASP and verify that this is in an RB-independent, non-canonical pathway.

Our data suggest that melanoma, pancreatic adenocarcinoma, and colorectal adenocarcinoma tumors with low *CDKN2A* expression have decreased SASP factor expression (Fig. 4). Importantly, our study shows that low *CDKN2A* expression correlates with downregulation of a distinctive SASP profile depending on the tissue of origin (Fig. 4B). This observation is consistent with previous studies suggesting that the SASP composition is temporally dynamic and context-and senescence inducer-dependent [53, 54, 87]. Characterization of the different SASP profiles and their unique dynamics will be critical not only to assess the senescent cell burden, but also to develop specific and personalized senescence-and SASP-targeted therapies. Since suppression of p16 can lead to senescence bypass and promote tumorigenesis [8–10, 12, 70, 71], obtaining profiles of the SASP factors related to this process may help treat the ~50% of all human tumors with low p16 expression [58].

It is well-established that the SASP has pleiotropic, context-dependent effects that both promote tumor progression, but also enhance anti-tumor immunity [reviewed in [88]]. For example, IL6 promotes chronic inflammation and tumorigenesis [89]. However, recent studies suggest that IL6 can also enhance anti-tumor immunity by resculpting T cell-mediated immune responses [89]. Likewise, other SASP factors, such as IL1a, IL1b, and TNF have this dual role where they can both promote inflammation and tumorigenesis or impair malignant transformation of benign nevi [90]. Using data from TCGA, we found that melanoma, pancreatic adenocarcinoma, and colorectal adenocarcinoma patients with low *CDKN2A* expression have a decreased SASP signature as well as decreased Antigen Processing Cross Presentation signaling [i.e., the ability of antigen presenting cells to present antigens on major histocompatibility complexes (MHCs) to T-cells]. Consistent with our observation, it has been shown that OIS primary human melanocytes upregulate the MHC class II apparatus to induce T-cell proliferation and that melanoma patients that sustain this feature have favorable disease outcome [91]. Additionally, suppression of p16 activity has been associated with immune deserts, immune escape, and low cytolytic activity in melanoma and pancreatic adenocarcinoma [92–94]. Thus, it is possible that in the context of certain tumor types such as those studied here, the decreased expression of SASP factors observed upon p16 knockdown or in *CDKN2A*-low patients may contribute to abrogation of senescence surveillance by immune cells [47, 95], thereby promoting tumorigenesis. Future experiments will determine whether suppression of p16 leads to decreased immune surveillance and the mechanism whereby this occurs.

Although loss of p16 is one of the most common events in cancer (~50% of all human cancers), there are currently no approved targeted therapies [80]. Additionally, we and others have shown that suppression of p16 has roles outside the cell cycle that would not be affected with current therapies undergoing clinical trials targeting CDK4/6 [80]. Therefore, finding downstream targetable pathways may be beneficial for this large subset of patients. For instance, we previously showed that inhibition of nucleotide metabolism through suppression of mTORC1 or the pentose phosphate pathway enzyme Ribose 5-Phosphate Isomerase A (RPIA) induces senescence specifically in p16-low cancers [8]. Here we found that suppression of p16 leads to decreased SASP expression. Therefore, induction of senescence in p16-low cancers may be a valuable strategy to inhibit the cell cycle while not activating the potential deleterious effects of the SASP.

In summary, we found that suppression of p16 decreases expression of SASP genes *IL6* and *CXCL8*, which cannot be explained by inhibition of senescence. We found that this phenomenon also occurs in p16-wildtype tumor cells upon p16 knockdown, and there is a decrease in the SASP gene signature in multiple tumor types that is associated with low p16 expression. Understanding whether p16 regulates SASP expression is critical to understand the complex relationship between cellular senescence, the immune system, and the cell cycle, three key players in cancer regulation.

## Methods

### Cell lines

Normal diploid IMR90 human fibroblasts were obtained from ATCC (CCL-186) and cultured according to the ATCC protocol in DMEM (Corning, cat#10-017-CV) supplemented with 5% FBS (VWR, cat#97068-085), L-glutamine (Corning, cat#25-015-CI), non-essential amino acids (Corning, cat#25-025-CI), sodium pyruvate (Corning, cat#25-000-CI), and sodium bicarbonate (Corning, cat#25-035-CI). Cells were cultured under physiological oxygen conditions (2% O2) and 5% CO2. Normal skin fibroblasts derived from a melanoma patient Hs 895.Sk were obtained from ATCC (CRL-7636) and cultured in DMEM (Corning, cat#10-013-CV) supplemented with 10% FBS (VWR, cat#97068-085). Experiments were performed on IMR90 between population doubling #25-35 and in Hs 895.Sk between population doubling #4-10. Melanoma cell lines SKMel28, Hs 600.T and RPMI-7951, were obtained from ATCC (HTB-72, CRL-7368, and HTB-66, respectively). SKMel28 and Hs 600.T as well as the lentiviral and retroviral packaging cells (293FT and Phoenix, respectively) were cultured in DMEM (Corning, cat#10-013-CV) supplemented with 10% FBS, while RPMI-7951 were cultured in EMEM (ATCC, cat#30-2003) supplemented with 10% FBS. Hs 895.Sk and cancer cell lines were cultured under 5% CO2. All cell lines were cultured in MycoZap (Lonza, cat#VZA-2032) and were tested for mycoplasma every two months as described in [96]. All tumor cell lines express wildtype *CDKN2A* according to TCGA [97, 98].

### Lentiviral and retroviral packaging and infection

pBABE BRAF^V600E^ (Addgene cat#15269), pBABE HRAS^G12V^ (Addgene cat#9051), and pBABE empty control (Addgene cat#1764) vectors were packaged into retroviral particles using the BBS/calcium chloride method as previously described in [8]. pLKO.1-shp16 (Sigma-Aldrich, TRCN0000010482) and pLKO.1-shGFP control (Addgene, cat#30323) vectors were packaged using the ViraPower Kit (Invitrogen, cat# K497500). Experimental timelines for IMR90 and Hs 895.Sk are delineated in Fig. 1A&2A. Briefly, cells were infected with pBABE control, pBABE BRAF^V600E^, or pBABE HRAS^G12V^ retroviral particles, and 24 hours later cells were infected with a second round of corresponding retroviral particles. Cells were infected with pLKO.1-shp16 or pLKO.1-shGFP when indicated in Fig. 1A&2A. Cells were selected with 1µg/mL puromycin for single infections or 3µg/mL for double infections until the end of the experimental procedure.

### RT-qPCR

Total RNA was extracted from cells with Trizol (Ambion, cat#15596018) and DNase treated, cleaned, and concentrated using Zymo columns (Zymo Research, cat#R1013) following the manufacturer’s instructions. Optical density values for RNA were measured using NanoDrop One (Thermo Scientific) to confirm an A260 and A280 ratios above 1.9. Relative expression of *IL6*, *CXCL8*, and *LMNB1* were analyzed using the QuantStudio 3 Real-Time PCR System (Thermo Fisher Scientific) with clear 96-well plates (Greiner Bio-One, cat#652240). Primers were designed using the Integrated DNA Technologies (IDT) web tool (*IL6*: forward 5’-GCCCAGCTATGAACTCCTTCT −3’ and reverse 5’-GAAGGCAGCAGGCAACAC −3’, *CXCL8*: forward 5’-AGACAGCAGAGCACACAAGC −3’ and reverse 5’-ATGGTTCCTTCCGGTGGT −3’, *LMNB1*: forward 5’-GAGAAGGCTCTGCACTGTATAC −3’ and reverse 5’-TGGAGTGGTTGTTGAGGAAG −3’, *MRPL19*: forward 5’-CAGTTTCTGGGGATTTGCAT −3’ and reverse 5’-TATTCAGGAAGGGCATCTCG −3’, *PSMC4*: forward 5’-TGTTGGCAAAGGCGGTGGCA −3’ and reverse 5’-TCTCTTGGTGGCGATGGCAT −3’ and *PUM1*: forward 5’-CGGTCGTCCTGAGGATAAAA −3’ and reverse 5’-CGTACGTGAGGCGTGAGTAA −3’). A total of 25ng of RNA was used for One-Step qPCR (Quanta BioSciences, cat# 95089-200) following the manufacturer’s instructions in a final volume of 10uL. Conditions for amplification were: 10 min at 48°C, 5 min at 95°C, 40 cycles of 10 sec at 95°C and 7 sec at 62°C. The assay ended with a melting curve program: 15 sec at 95°C, 1 min at 70°C, then ramping to 95°C while continuously monitoring fluorescence. Each sample was assessed in triplicate. Relative quantification was determined to multiple reference genes (*MRPL19*, *PSMC4* and *PUM1*) to ensure reproducibility using the delta-delta CT method.

### Western blotting

Cell lysates were collected in 1X sample buffer (2% SDS, 10% glycerol, 0.01% bromophenol blue, 62.5mM Tris, pH=6.8, 0.1M DTT) and boiled (10 min, at 95°C). Protein concentration was determined using Bradford assay (Bio-Rad, cat#5000006). Proteins were resolved using SDS-PAGE gels and transferred to nitrocellulose membranes (GE Healthcare Life Sciences, cat#10600001) as previously described in [8]. Antibodies used include: anti-BRAF (Santa Cruz Biotechnology, cat#sc-5284, 1/1000 dilution), anti-RAS (BD Sciences, cat#610001, 1/1000 dilution), anti-p16 (Abcam, cat#ab108349, 1/1000 dilution), anti-vinculin (Sigma-Aldrich cat#V9131, 1/1000 dilution), β-Actin (Sigma-Aldrich, cat#A1978, 1/10000 dilution), anti-mouse HRP (Cell Signaling Technology, cat#cst7076, 1/10000 dilution), and anti-rabbit HRP (Cell Signaling Technology, cat#cst7074, 1/5000 dilution).

### Differential expression analysis

Preprocessed and processed RNA-Seq data from skin cutaneous melanoma (SKCM), pancreatic adenocarcinoma (PAAD), and colorectal adenocarcinoma (COADREAD) TCGA data were downloaded from BROAD GDAC Firehose on June 22, 2020 [Broad Institute TCGA Genome Data Analysis Center (2016): Firehose 2016_01_28 run. Broad Institute of MIT and Harvard. doi:10.7908/C11G0KM9)]. Processed rnaseqv2 files containing normalized RSEM expression values for each gene in each patient were used to determine the first and third quartile of *CDKN2A* expression for each tumor type separately (Table 1). Quartile values were used to classify patients into low-p16 expression (*CDKN2A* expression ≤ Q_1_) and high-p16 expression (*CDKN2A* expression ≥ Q_3_). Differential expression analysis between p16-low and p16-hihg patients for each tumor type was performed using the preprocessed raw-counts files in R-CRAN (R-3.6.3) and the DESeq2 package.

### Gene Set Enrichment Analysis (GSEA)

Genes were ranked according to the fold-change and p-value obtained in the differential expression analysis as follows: −log_10_(p-value) × sign (log_2_ fold change). Pre-ranked files were built for each tumor type separately in R-CRAN (R-3.6.3) and used to run pre-ranked GSEA (javaGSEA desktop application) for Hallmarks, KEGG and Reactome under predefined parameters (1000 permutations, weighted enrichment statistic, excluding sets larger than 500 and smaller than 15 and using meandiv normalization mode, there were no repeated genes thus collapse mode was not used). Following GSEA documentation, indications terms were considered significant when the FDR adjusted p-value (q-value) was <0.25 (http://software.broadinstitute.org/gsea/index.jsp).

### Statistical Analysis

GraphPad Prism version 7.0 was used to perform statistical analysis. The level of significance between two groups was assessed with unpaired t test. For data sets with more than two groups, one-way ANOVA followed by Tukey’s post hoc test was applied. P-values< 0.05 were considered significant. Pearson correlation test in GraphPad prism version 7.0 was used to assess the correlation between *LMNB1* and *CDKN2A*. Venny 2.0 tool was used to obtain common downregulated genes among tumor types (Fig. 4B and **Table S3**) as well as common GSEA terms (**Table S4**).

## Acknowledgments

We thank Erika Dahl and Drs. Chi-Wei Chen and Naveen Kumar Tangudu for critical reading and editing of this manuscript. We also thank Dr. Gary Nolan from Stanford University for providing with Phoenix packaging cells.

## Supplemental Materials

**Table S1:** 232 total unique SASP genes including soluble factors and exosomes obtained from (Basisty et al., 2020)

**Table S2:** Differential expression analysis of *CDKN2A*-low vs. *CDKN2A*-high tumors obtained from TCGA

**Table S3:** Cross-comparison of significant downregulated SASP among the three tumor types

**Table S4:** Cross-comparison of the Gene Set Enrichment Analysis output with negative NES for the tumor types

